# A behavioral syndrome linking boldness and flexibility facilitates invasion success in sticklebacks

**DOI:** 10.1101/2021.02.06.430052

**Authors:** Miles K. Bensky, Alison M. Bell

**Affiliations:** Program in Ecology, Evolution, and Conservation Biology, School of Integrative Biology, University of Illinois, 505 S. Goodwin Ave., Urbana, IL 61801; Department of Evolution, Ecology and Behavior, Carl R. Woese Institute for Genomic Biology, Neuroscience Program, University of Illinois, Urbana, Illinois 61801

**Keywords:** plasticity, persistence, novel object, intraspecific variation, marine, freshwater, convergent, rapid adaptation

## Abstract

For a species to expand its range, it needs to be good at dispersing and also capable of exploiting resources and adapting to different environments. Therefore, behavioral and cognitive traits could play key roles in facilitating invasion success. Here, we show that dispersing sticklebacks are bold, while sticklebacks that have recently established in a new region are flexible. Moreover, boldness and flexibility are negatively correlated with one another at the individual, family and population levels. Multiple lines of evidence suggest that the divergence in boldness and flexibility is likely to be evolutionary in origin. If boldness is favored in invaders during the initial dispersal stage, while flexibility is favored in recent immigrants during the establishment stage, then the link between boldness and flexibility could generate positive correlations between successes during both dispersal and establishment, and therefore play a key role in facilitating colonization success in sticklebacks and other organisms.

## INTRODUCTION

Understanding the factors that allow a species to expand its range and adapt to changing habitats is increasingly important in the face of anthropogenic change. Natural biological invasions can reveal how and why certain organisms can excel in response to novel selection pressures (Whitney & Gabler 2008). In addition to the importance of propagule pressure (Lockwood *et al.* 2005), stochasticity, and the opening of new niches at the edge of species boundaries, there is growing evidence that particular traits (e.g., r-selected life histories (Capellini *et al.* 2015), habitat breadth (Blackburn *et al.* 2009), large brains/cognitive abilities (Sol *et al.* 2005)) might promote biological invasions.

Behavioral and cognitive traits are likely to play an important role in allowing animals to move into and become established in new environments (Sih *et al.* 2010; Wright *et al.* 2010; Hoset *et al.* 2011; Chapple *et al.* 2012). For example, behavioral plasticity allows animals to rapidly adjust their phenotype in response to changes in environmental cues (Wolf *et al.* 2008; Coppens *et al.* 2010; Sih *et al.* 2012). Cognitive processes – how animals perceive, process and retain information about their environment and then use that information to make decisions (Shettleworth 2010) – may play an especially important role during biological invasions because they influence the ability of animals to enter new habitats, exploit new niches, become established and spread (Wright *et al.* 2010; Griffin *et al.* 2016; Szabo *et al.* 2020). For example, recent immigrants have to be willing to approach and interact with novel stimuli in order to gain information about their new environment (i.e., neophilia (Mettke-Hofmann *et al.* 2009)). Additionally, previously successful behavioral patterns may no longer be successful in new environments, so immigrants need to be able to stop persisting on ineffective responses and flexible enough to attempt new approaches (Griffin *et al.* 2016).

Successfully colonizing a new environment can be broken down into discrete stages, e.g., dispersal, colonization, establishment and spread (Chapple *et al.* 2012), and invasion success likely relies on different behavioral and cognitive traits in each stage (Sih *et al.* 2012). For example, dispersers need to be bold, while immigrants can succeed in a new environment when they are willing to investigate novel stimuli and when they are able to quickly inhibit old and ineffective behaviors, i.e., if they are flexible (Sih *et al.* 2012). If boldness, neophilia and behavioral flexibility are no longer beneficial or even costly once a population becomes established in a new environment, they may be lost (Wright *et al.* 2010; Herczeg *et al.* 2020). Numerous studies have documented differences in behavioral traits between invading and established populations, e.g., (Martin & Fitzgerald 2005; Pintor *et al.* 2008; Damas-Moreira *et al.* 2019; Cohen *et al.* 2020), and a handful of studies have shown that they have a heritable basis, e.g., (Sargent & Lodge 2014; Gruber *et al.* 2017). If there is an underlying genetic basis to the behavioral and cognitive traits that facilitate range expansions or biological invasions, then we might expect those traits to evolve over the course of an invasion. According to this hypothesis, behavioral and cognitive traits important for invasion should vary in a systematic way between dispersing populations compared to newly-derived and well-established populations when they are reared in a common garden. Moreover, if invasion success requires different behavioral and cognitive traits in the different invasion stages, then mechanisms that package these traits together could be key to the success of invasive species (Sih *et al.* 2012). For example, traits important for the dispersal and establishment stages are coupled together in western bluebirds, and this facilitates the expansion of their range (Duckworth & Badyaev 2007).

Threespined sticklebacks (*Gasterosteus aculeatus*) are a model system for studying trait evolution during biological invasions. Throughout their evolutionary history, marine sticklebacks have repeatedly colonized freshwater environments, rapidly adapted to them and diversified (Stuart *et al.*). Sticklebacks can also spread and have dramatic impacts on freshwater communities (Eklöf *et al.* 2020; Gugele *et al.* 2020). Work on this system has primarily focused on a suite of morphological and physiological traits that repeatedly evolves when marine sticklebacks invade freshwater habitats (Bell *et al.* 2004; Colosimo *et al.* 2005), with evidence that haplotypes containing a set of coadapted alleles are maintained at low frequency in the ocean and are repeatedly tapped during adaptation to freshwater (Bassham *et al.* 2018). However, little is known about the behavioral and cognitive mechanisms that may facilitate the invasion of sticklebacks into new habitats and which may evolve once a population becomes established (Foster *et al.* 2015).

We took advantage of the repeated invasion of freshwater habitats by sticklebacks by comparing marine stickleback to sticklebacks from recently-derived and well-established freshwater populations in Alaska, with replicate populations of each type. For each population, we reared clutches of field-fertilized eggs in a controlled laboratory environment and scored individuals for three informative and repeatable behavioral traits: boldness, behavioral flexibility, and neophilia (Bensky & Bell 2020). We hypothesized that boldness is favored in dispersers, that neophilia and behavioral flexibility are favored in newly-arrived immigrants, and that these traits may be subject to relaxed selection and possibly lost once a population becomes established. The replicate populations allow us to assess the generality of the patterns and comparing the newly-established to well-established populations provides insight into whether different traits are favored during initial establishment versus population persistence (Merwin 2019). Rearing animals under common-garden conditions in the lab also allowed us to determine if behavioral variation within and among populations likely reflect evolved, genetically based differences.

## MATERIALS AND METHODS

Adult sticklebacks were collected via minnow traps from six populations ranging from the Matanuska-Susitna Valley to the Kenai Peninsula of Alaska (Figure S1; Table S1) during June 2017. Two populations (Rabbit Slough and Resurrection Bay) occur in the ancestral marine environment, while the remaining four populations occur in freshwater. Two of the freshwater populations (Big Beaver and Cornelius, hereafter referred to as “well-established”) are derived from natural colonization events that presumably occurred hundreds to thousands of years ago, after the last glacial maximum, while the other two freshwater populations are recently derived via natural recolonization (Loberg: 28-34 years prior to collection; (Bell *et al.* 2004)) or experimental seeding (Cheney: 8 years prior to collection; (Bell 2016)). As is typical for sticklebacks, the marine populations are only weakly genetically differentiated from each other (F_ST_ = 0.0076; (Hohenlohe *et al.* 2010)) while the freshwater populations are more strongly genetically differentiated (M. Bell, K Veeramah, personal communication).

Eggs were fertilized in the field following previously established protocols (see (Wund *et al.* 2012)). Two to three days post fertilization, the eggs were transferred to 50 mL canonical tubes and shipped overnight in coolers filled with ice packs to the University of Illinois Urbana-Champaign where they were raised in common garden conditions in the lab. Artificial incubation controls for environmental paternal effects due to receiving paternal care therefore it is likely that phenotypic differences among the lab-reared populations reflect heritable differences (although environmental maternal effects could also contribute).

Clutches were reared in separate tanks (9.5L 32 × 21 × 19 cm) where the embryos were incubated in a cup with a mesh bottom and placed over an air bubbler. Fish were kept at 60°F with an even light cycle (12L:12D) for the entirety of the experiment. All families were kept on one of two recirculating flow-through water racks, which consisted of a series of particular, biological, and IV filters and had three different shelves (Aquaneering, San Diego, USA). 10% of each tank’s water was replaced each day. Family tank position was pseudo-randomly assigned so that all populations were evenly distributed across both racks and the three levels of shelves. Importantly, we elected to rear the fish and measure their behavior in freshwater (~5ppt), thereby simulating the conditions that marine sticklebacks encounter when they move into freshwater. Because the marine populations studied here are naturally anadromous (Rabbit Slough: (Bell 2016); Resurrection Bay: R. King, pers. comm.), i.e., they spawn in fresh/brackish waters, their early offspring development typically occurs under low salinity.

### Behavioral assays

#### Summary of the behavioral assays

Neophilia was measured as response to a novel object. Boldness was measured as latency to emerge from a refuge, a reliable and widely-used behavioral assay in fishes (Wilson & Godin 2009; Pearish *et al.* 2013). Boldness was quantified at the individual level as average latency to emerge across four independent trials. Behavioral flexibility was measured in a barrier task: after pretraining individuals to expect a food reward upon emergence from a refuge, individuals were confronted by a transparent barrier that they had to swim around in order to get the food reward. Individuals that continue to follow the prepotent search pattern established during training spend relatively more time at the point of the barrier closest to the food reward (“barrier apex”), which we interpret as relatively low flexibility. In contrast, individuals that quickly break away from the previously-established behavior pattern spend relatively little time spent at the apex of the barrier, which we interpret as relatively high flexibility. A previous study in sticklebacks found that time at the apex of the barrier predicts reversal learning performance (Bensky & Bell 2020), another common metric of behavioral flexibility.

#### Detailed experimental methods

18 observation tanks (36L × 33W × 24H cm) were used for behavioral testing. When the fish were approximately eight months of age (approximately 40 mm standard length), the testing phase of the experiment began. Behavioral assays were carried out over the course of 5 months. Families, sexes and populations were measured in a pseudorandomized order, such that male and female offspring from different families and from different populations were measured in the same block. Individuals were randomly selected from each family, and their weight and length were measured. During the testing phase of the experiment, fish were only fed during the behavioral tests to help maintain motivation.

### Acclimation phase

In order to ensure that an individual had acclimated to their home tank and was motivated to eat during the behavioral tests, the individual was presented with food via a petri dish at the center of their home tank, and the individual had to eat the food within 10 minutes on three consecutive days in order to proceed to the next step. On average, it took 5.1 days for fish to meet this criterion (range = 3 to 17 days).

### Novel object test (neophilia)

Individuals’ response to a novel object (toy lion; 10L × 7H cm; TERRA by Battat, Montreal, Canada) was recorded the day after the fish met criterion in the acclimation phase. The toy lion was selected as a novel object because the fish had no prior experience with this object, there was no presumed evolutionary history with the object’s shape, and it was made up of neutral colors.

The individual’s behavior was measured for five minutes after their first approach to the novel object (i.e., first time within one body length of the novel object and oriented directly towards it). We interpret more time spent near and oriented towards the novel object as greater time investigating the object (i.e., higher neophilia).

### Latency to emerge (boldness)

For each individual, we recorded their latency to emerge from a refuge on their first four training trials for the barrier task (described below) and used the average of those measures as a proxy for boldness (Movie S1). Latency to emerge was repeatable across the four trials within each population (r=0.56-0.72, Table S4).

### Barrier detour task (flexibility)

Pretraining for the barrier detour task started on the same day the novel object test was completed. The goals of pretraining were to train the individual to learn that there would consistently be a food reward in the middle of the tank, establish food motivation in this context, and create a prepotent response of leaving a shelter to directly approach and eat the food reward.

During pretraining, individuals were trained for one session per day, with each session comprising four trials. To start the trial, the observer removed a cork from the side of the shelter and the fish was given ten minutes to exit. Upon exiting the fish was allowed five minutes to eat the worm.

After eating the worm, the fish was placed back into the shelter in preparation for the next trial. If the fish did not emerge from the shelter within ten minutes after the cork was pulled or eat within five minutes after emergence, the observer recorded the maximum times for these behaviors, removed the food reward and gently poured the fish out of the shelter if necessary. Latency to emerge from a refuge during the first pretraining session was used as a measure of boldness (see above).

Training for the barrier task was criterion based. In order to move on to the barrier task following pretraining, the individual had to emerge from the shelter within 10 minutes and directly approach and eat the food reward within five seconds on three out of the four trials. The one failed attempt could not be on the fourth trial; this requirement ensured that the fish would be motivated throughout the four trials. Fish were given a maximum of four days to reach criterion. Fifteen fish did not meet criterion (Big Beaver: 7, Cornelius: 1, Loberg: 1, Cheney: 3, Rabbit Slough: 2, Resurrection Bay: 1).

Once an individual met criterion, the individual moved on to the barrier detour task the following day. This task also comprised four trials. The first two trials were exactly the same as the pretraining trials in order to reinforce the direct search pattern. On the third trial a transparent semi-circular barrier was placed between the shelter and food reward. The opening into the barrier area was positioned directly in front of the entrance to the shelter. After removing the cork the individual was allowed 30 minutes to emerge from the shelter, navigate around the barrier and eat the food reward (Movie S2). The observer recorded the duration of the first bout (no break in contract longer than five seconds) at the apex of the barrier. In order to confirm that the fish that spent little time at the barrier apex during the third trial were still motivated to eat, the fish’s behavior was observed for a fourth trial during which no barrier was present.

Altogether, a total of n=262 individuals from n=8-11 families per population (n=2-7 full sibs per family) completed the novel object test and boldness assay. A total of n=247 individuals from n=8-11 families per population (n=1-4 full sibs per family) completed the barrier task. The experiments were approved by the Institutional Animal Care and Use Committee (IACUC) of the University of Illinois Urbana-Champaign (IACUC protocol #15077).

### Statistical analysis

R 3.5.3 (http://www.r-project.org/) was used for statistical analyses. Positively-skewed variables were log-transformed to improve normality, model residuals were also visually inspected for deviations from normality.

We used linear mixed models (Team 2013)(package = “lme4”; function = “lmer”) to examine the behavioral data in each assay separately. We created models in which population and sex were included as fixed factors, and body length was included as a covariate. FamilyID nested within population was included as a random variable. The statistical significance of the effect of FamilyID was assessed by AIC (Akaike 1973), i.e., by comparing models with and without the effect of FamilyID (R Core Team 2016; package = “care”; function = “anova”). We infer that a trait has an underlying heritable basis when the trait differs among populations (because they were reared in a common garden), and/or when FamilyID improves model fit. To further explore the heritable basis to these traits we also computed broad-sense heritabilities and genetic correlations within each population following (Hadfield 2010; Wilson *et al.* 2010). We used weakly informative inverse-gamma priors for the ‘residual’ and ‘genetic’ effects (by setting the MCMCglmm parameters *V*◻=◻1, nu◻=◻0.002). In order to test for genetic correlations, a 2×2 covariance matrix was specified with the degree of belief parameter set to n=1.002. The raw positively-skewed duration data for all the behavioral measures were used for this analysis, and thus all measures were rounded to the nearest integer and a Poisson distribution was used. The posterior distribution was sampled every 3000 times (thinning interval), following a burn-in period of 100◻000 iterations with a total run of 10◻000◻000 iterations. The relatively modest number of families and full sibs within each family resulted in large confidence intervals around the estimates therefore we interpret those results with some caution.

To examine the structure of the phenotypic variation both among and within populations, we examined correlations between traits at the individual, family and population level.

See SI Methods for more details.

## RESULTS

### Sticklebacks from dispersing populations were bold

There was significant variation among populations in boldness (latency to emerge from a refuge (F_5,59_ = 3.30, p = 0.01, Table S1, Figure 1)). Sticklebacks from dispersing populations (marine) were more bold than fish from well-established freshwater populations (z = 2.745, n = 262, p = 0.006). The two recently-derived populations provide insight into the rate at which boldness diverged from the ancestral marine behavioral type. Sticklebacks from Cheney Lake, a population that was established eight years prior to collection, resembled marine populations and differed from the well-established freshwater populations (Cheney vs. well-established: z = −2.497, n = 262, p = 0.034). In contrast, sticklebacks from Loberg Lake, a population that was established 28-34 years prior to collection, more closely resembled the well-established freshwater populations (Loberg Lake vs marine: z = 2.429, n = 262, p = 0.040). After correcting for multiple comparisons, there was a trend for sticklebacks from Cheney Lake to emerge faster compared to sticklebacks from Loberg Lake (z = −2.249, n = 91, p = 0.064).

**Figure 1.**
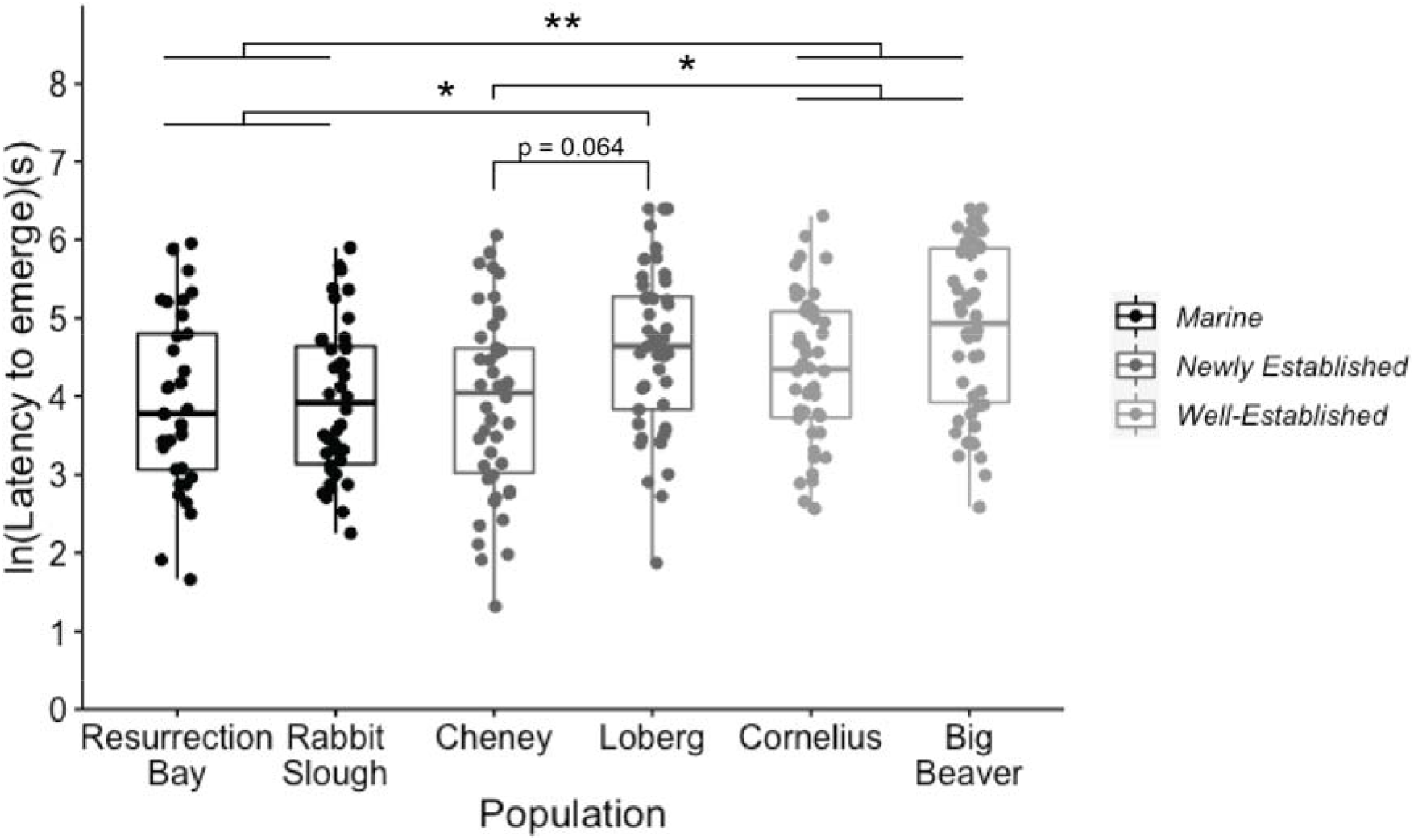
Variation among populations in boldness (latency to emerge from a refuge). Smaller values indicate greater boldness. Figure shows boxplots with individual data points superimposed.

There was significant variation in boldness among families within populations (FamilyID improved model fit (X^2^ = 9.394, df = 1, p = 0.002; AIC_with_ = 774.50; AIC_without_ = 781.89)). Estimates of broad sense heritability of boldness in the six populations ranged from 0.1 to 0.75 (average = 0.48) and was significantly different from zero in the Big Beaver, Cornelius and Resurrection Bay populations (Table S2).

### Sticklebacks from established populations were flexible

There was significant variation among populations in behavioral flexibility (time at the apex of a barrier (F_5,58_ = 3.495, p = 0.008, Table S1, Figure 2)). Sticklebacks from the well-established populations were more flexible (spent less time at the apex of a barrier), compared to sticklebacks from the dispersing marine populations (z = −2.175, n = 158, p = 0.030). As was the case for boldness, the two recently-derived populations show different patterns: flexibility in Cheney Lake – the most recently established freshwater population – resembled flexibility in the dispersing populations, while flexibility in Loberg Lake – the freshwater population that was established 28-34 years prior to collection – resembled flexibility in the well-established freshwater populations (Cheney Lake vs Loberg Lake: z = 2.821, n = 87, p = 0.013, Cheney Lake vs well-established: z = 3.373, n = 131, p = 0.002).

**Figure 2.**
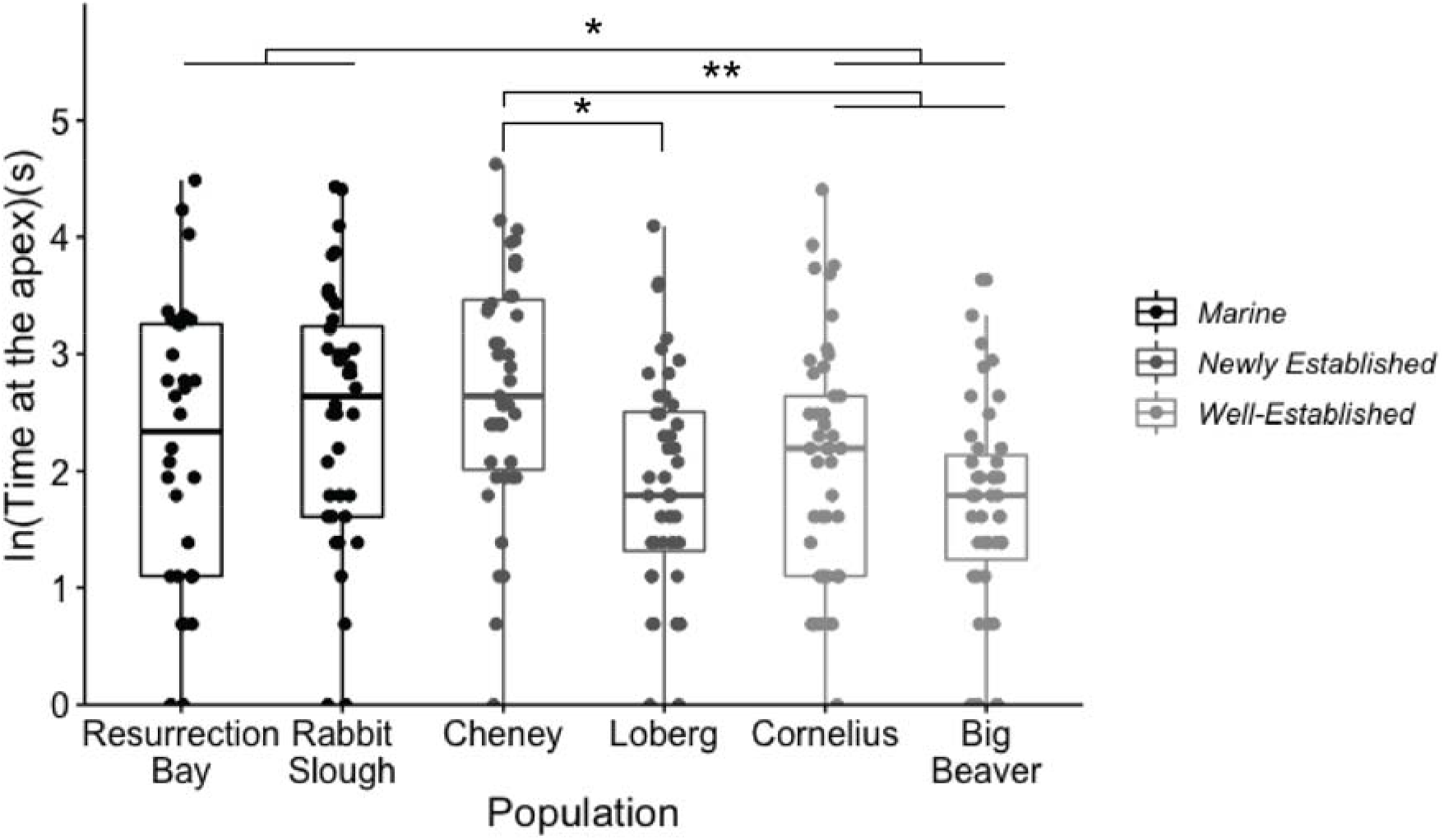
Variation among populations in flexibility (time at the apex of the barrier). Smaller values indicate greater flexibility, i.e., less time persisting on a previously-successful behavior pattern. Figure shows boxplots with individual data points superimposed.

There was significant variation among families within populations in flexibility (X^2^ = 7.350, df = 1, p = 0.007; AIC_with_ = 723.74; AIC_without_ = 729.09). Estimates of broad sense heritability of flexibility in the six populations ranged from 0.22 to 0.86 (average = 0.45) and was significantly different from zero in the Cornelius population (Table S2).

### Neophilia did not evolve systematically during the invasion

Neophilia (time near and oriented to a novel object) did not vary among populations or between the sexes (Table S1) and did not vary in a systematic manner among the different types of populations (Figure 3). Larger fish were less neophilic (β = −0.033, t = −2.631, df = 247.79, p=0.009, Figure S2, Table S1). We did not detect variation among families within populations in neophilia, as FamilyID did not significantly improve model fit (X^2^ = 2.369, df = 1, p = 0.124; AIC_with_ = 636.57; AIC_without_ = 636.94). Estimates of broad sense heritability of neophilia in the six populations ranged from 0.09 to 0.87 (average = 0.34) and was significantly different from zero in the Cornelius population (Table S2).

**Figure 3.**
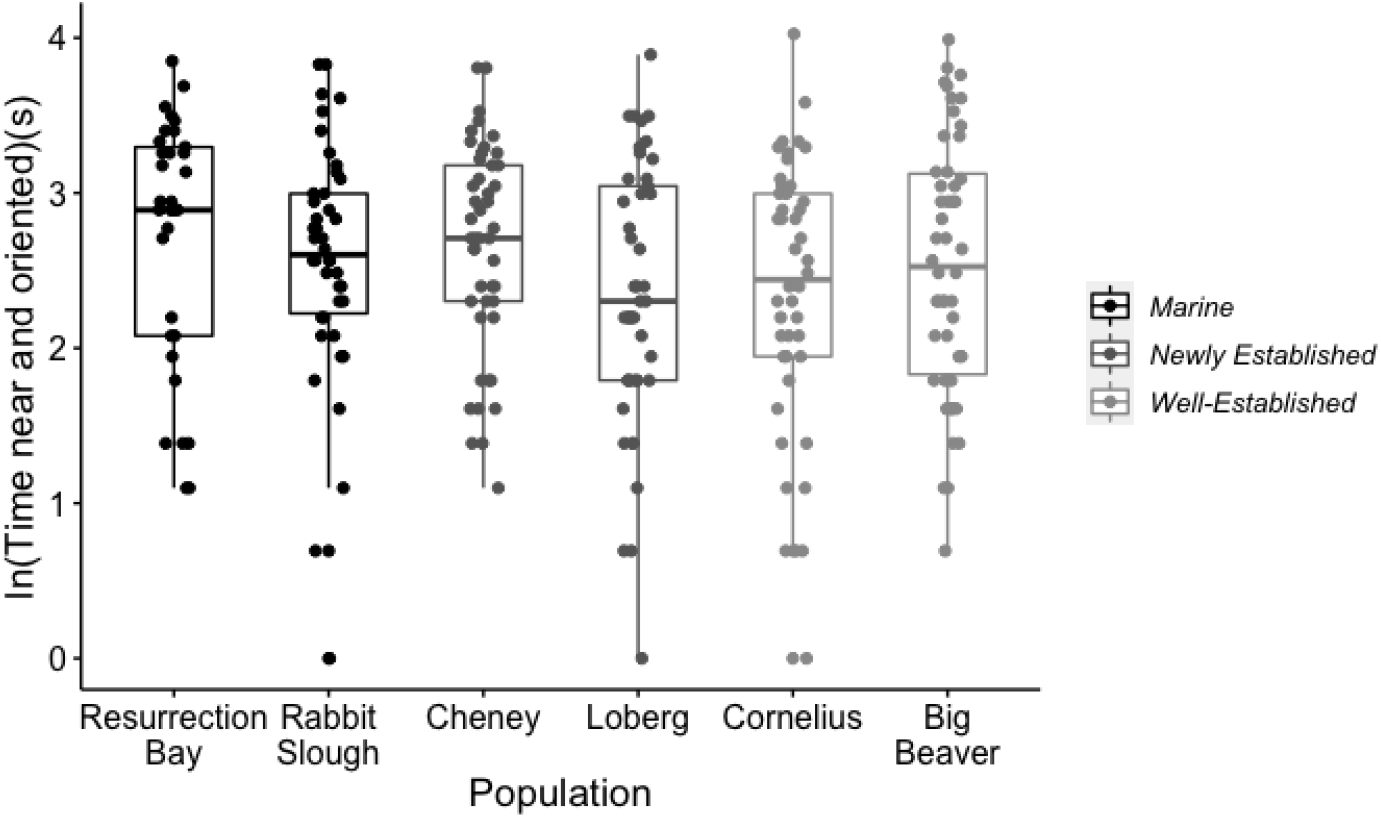
Variation among populations in neophilia (time near and oriented to the novel object). Figure shows boxplots with individual data points superimposed.

### A boldness-flexibility syndrome facilitates invasion success

Given that sticklebacks from the dispersing populations were bolder and less flexible compared to sticklebacks from established populations, we tested how boldness and flexibility were correlated with one another. Consistent with the pattern described above, individuals that were more bold (quickly emerged from the refuge) were less flexible (spent more time at the barrier apex; r = −0.522, n = 245, p<0.001, Table S3), and this pattern was also evident at the family (r = −0.688, n = 62, p<0.001) and population (r = −0.91, n = 6, p=0.01) levels (Figure 4). The average genetic correlation between boldness and flexibility within the six populations was r = −0.69 (range: −0.59 to −0.77) and was significantly different from zero in the Cornelius population (Table S3).

**Figure 4.**
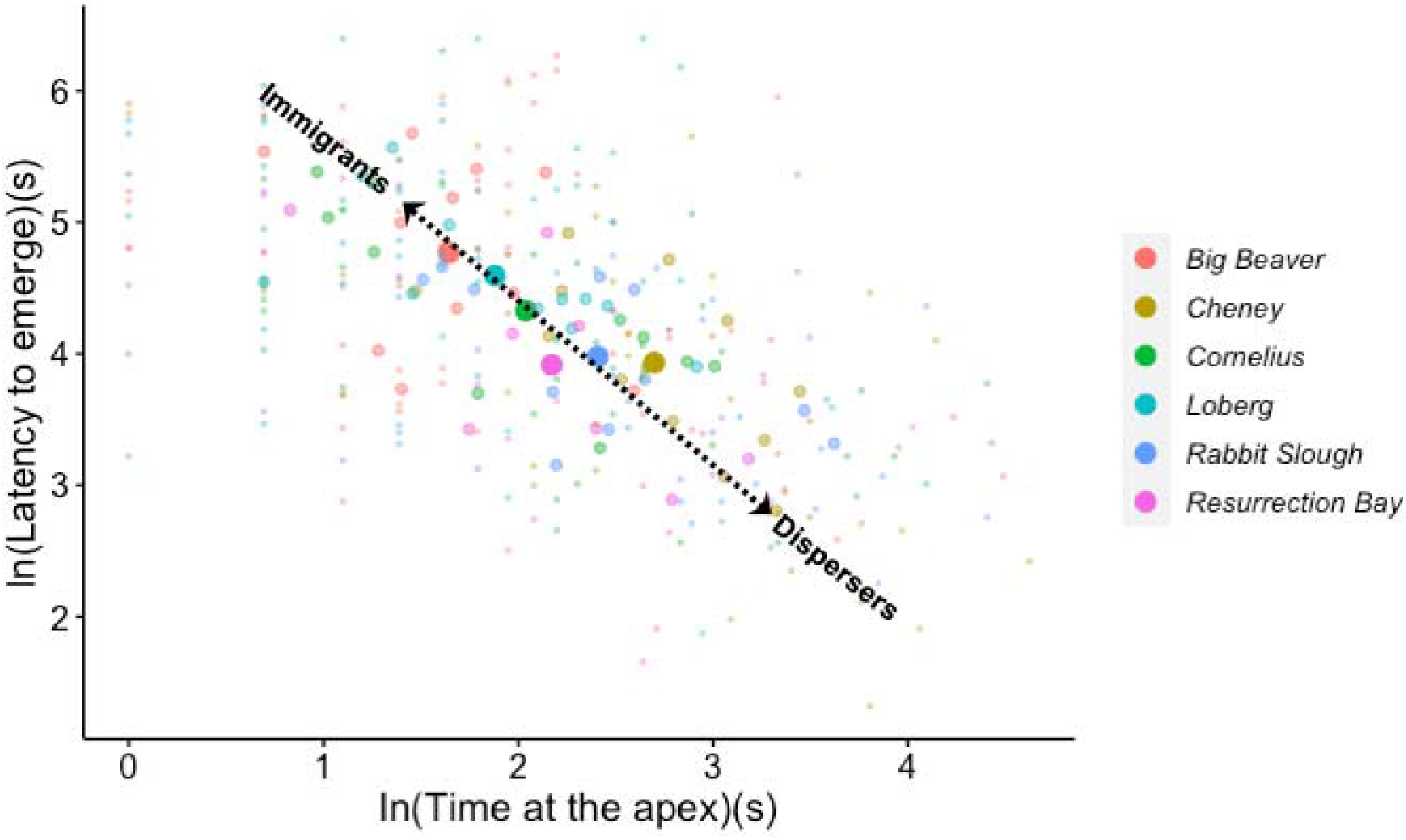
Relationship between flexibility (time at the barrier apex) and boldness (latency to emerge) within and among populations. Dispersers are relatively bold (emerge quickly) while immigrants are relatively flexible (spend less time at the barrier apex). Large circles represent populations means, medium circles show family means, small circles show individual data points. Circles are color coded by population. The line and text drawn on the figure are for visual purposes only.

## DISCUSSION

The possibility that behavioral and cognitive traits might facilitate and evolve during natural biological invasions is intriguing but difficult to study directly. Here, we took advantage of a model system for invasions to test the hypothesis that boldness is favored in dispersers, that neophilia and flexibility are favored in recently-arrived immigrants and that these traits are subject to relaxed selection and possibly lost once a population becomes established in a new environment. We found that boldness and flexibility evolve in a systematic way when marine sticklebacks colonize freshwater habitats. Specifically, the dispersing populations were bold, and well-established populations were flexible. These traits varied between the two recently established populations relative to time since establishment. Moreover, boldness and flexibility were negatively correlated with one another at the individual, family and population levels. Differences in boldness and flexibility were evident in a common garden environment, there was significant variation among families in both of these traits, and estimates of their heritability and the genetic correlation between them were relatively high for behavioral traits (Dochtermann *et al.* 2019). These lines of evidence suggest that there is a heritable component to the traits and that their divergence is likely to be evolutionary in origin.

If boldness is favored in invaders during the initial dispersal stage, while flexibility is favored in recent immigrants during the establishment stage, then a tight link between boldness and flexibility could generate positive correlations between successes during both the dispersal and establishment stage, and therefore play a key role in facilitating invasion success in this species (Sih *et al.* 2012). Selection favoring boldness in dispersers and flexibility in immigrants could cause the population to move along a ridge of high fitness whenever sticklebacks diverge from the ancestral marine behavioral type (Figure 4). An outstanding question is what maintains the extensive phenotypic and genetic variation in boldness and flexibility within the populations (Figures 1–4). Selection might not be strong enough to effectively purge the variation, and/or there may be ongoing gene flow between marine and freshwater habitats which maintains recessive alleles associated with behavioral flexibility at low frequency in marine populations (transporter hypothesis (Schluter & Conte 2009)), as appears to operate at the *Eda* locus (Jones *et al.* 2012) and surrounding genomic regions (Bassham *et al.* 2018).

We found no support for the hypothesis that traits which promote dispersal and early establishment in a new environment are lost once a population becomes well-established (Wright *et al.* 2010), i.e., no evidence that newly-derived populations were more bold, neophilic and flexible than well-established populations. One possible explanation for this pattern is that *assortative mating* within dispersing populations causes dispersing behavioral phenotypes to be maintained early in the establishment process, even in the presence of selection favoring more flexible phenotypes (Shine *et al.* 2011). The comparison between the two newly-derived populations (Cheney Lake and Loberg Lake, respectively) is also insightful; the two newly derived populations either tended to resemble the dispersing populations or the well-established populations. Cheney Lake was founded more recently than Loberg Lake (9 versus 28-34 years prior to this study); if it takes longer than 10 generations for behavioral and cognitive traits to diverge from the ancestral marine behavioral type then *time since establishment* could be important. Alternatively, or in addition, the phenotypic differences between sticklebacks from Loberg Lake and Cheney Lake could reflect differences in the way that the two lakes were colonized; Loberg Lake was naturally colonized, while sticklebacks were experimentally introduced to Cheney Lake. If particularly flexible individuals were more likely to disperse into Loberg Lake, but a random sample of behavioral types were artificially introduced into Cheney Lake (Cote *et al.* 2010; Edelaar & Bolnick 2012; Canestrelli *et al.* 2016), then *non-random dispersal* could be contributing to the rapid evolution of increased flexibility in Loberg Lake. Given evidence from the literature (Cote *et al.* 2010; Canestrelli *et al.* 2016) and from this study that bold individuals are more likely to disperse, and that boldness and flexibility are tightly *negatively* correlated with one another, this explanation seems unlikely. Further studies tracking how behavioral and cognitive traits change over time in the Cheney Lake population (and similar experimental lakes (e.g., Scout Lake)) could help discriminate between the assortative mating, time since establishment and non-random dispersal hypotheses.

We originally hypothesized that neophilia would be favored in recently-derived populations because seeking and/or being willing to investigate novel stimuli may help newly-arrived immigrants locate new habitats and discover novel resources, but we found no support for this hypothesis. One possible explanation for the failure to find systematic differences in neophilia among the populations is that neophilia may actually be disadvantageous in a new environment because it can expose animals to dangerous stimuli they have never encountered before. Another potential (nonexclusive) explanation based on our results is that neophilia may not evolve as readily because it may be less heritable (effect of FamilyID was nonsignificant, lower H^2^ estimate). Instead, neophilia may be more influenced by age or experience: in this study, smaller (and younger: r = −0.159, t = −2.595, n = 262, p=0.01) fish were more neophilic, which is consistent with other studies which have shown that novelty-seeking decreases with age (Stansfield & Kirstein 2006).

Sticklebacks are a powerful model system for understanding how and why certain traits repeatedly evolve whenever organisms invade new habitats. Accumulating evidence suggests that sticklebacks have evolved mechanisms for rapidly adapting to new environments with alleles conferring the freshwater-adapted phenotype maintained at low frequency in the ocean (Colosimo *et al.* 2005). But the role of behavior and cognition in facilitating evolutionary processes in this system has received less attention (Foster *et al.* 2015). Our results suggest that a behavioral mechanism – a behavioral syndrome linking boldness and flexibility together – can contribute to rapid adaptation in this and other organisms.

## ACKNOWLEDGEMENTS

We thank Matt Wund, Mike Bell, John Baker, and the members of the Foster Lab for help with field work. The Bell lab, Andy Suarez, Jason Keagy, Becky Fuller and Katy Heath provided valuable feedback. Financial support was provided by the National Science Foundation’s Integrative Graduate Education and Research Traineeship program and by the University of Illinois. This material is based upon work supported by the National Science Foundation under Grant No. IOS 1121980 and by the National Institutes of Health under award number 2R01GM082937-06A1.

