## Supplementary Information for "A behavioral syndrome linking boldness and flexibility facilitates invasion success in sticklebacks"

Miles K. Bensky

**This PDF file includes:**

Supplementary text

Figures S1 to S2

Tables S1 to S4

Legends for Movies S1 to S2

SI References

**Other supplementary materials for this manuscript include the following:**

Movies S1 to S2

Supplementary Information Text

**Supplementary methods.** Embryos were generated via artificial fertilization and incubation. To collect sperm, males from each population were euthanized using an overdose of buffered tricaine methanesolfonate (MS-222), and their testes were immediately dissected and macerated. Eggs were then gently extruded from gravid females from the same population into a petri dish and the macerated testes were pipetted over the eggs to fertilize them. Distilled water with 6 ppt Instant Ocean ® was used to repeatedly rinse the newly fertilized clutches before being stored in that solution in the petri dish. The clutches were inspected daily for proper development; dead embryos and unfertilized eggs were removed, and the water was changed.

Once the embryos arrived at the University of Illinois Urbana-Champaign, each clutch was transferred into a plastic cup with a mesh bottom and clipped to the side of a tank (one clutch per tank; (9.5L 32 x 21 x 19 cm)). A bubbler was placed under the cup to provide aeration. Tanks used for rearing embryos were lined with gravel and had a refuge (artificial plant). Embryos were checked daily and dead embryos were removed. Upon hatching (8-13 days post fertilization) fry were fed brine shrimp daily. Once the eggs had hatched the mesh cup and bubbler were removed.

At approximately two months of age, the fish were gradually introduced to a mixed diet of frozen bloodworms, frozen brine shrimp, and frozen Mysis shrimp and were fed ad lib once a day. To prevent overcrowding, families were culled to a maximum of 30 fish per tank at two months, and to a maximum of 15 fish at approximately six months. Throughout the experiment each family was maintained in its own tank. Therefore, tank effects could potentially contribute to differences among families. Tanks from each population were evenly distributed around the fish room in an attempt to control for location effects.

Tissue for DNA was collected by swabbing the side of each fish with a sterile cotton swab to collect DNA for non-invasively determining the sex of each fish with a genetic marker (Peichel *et al.* 2004). Total genomic DNA was extracted using the DNEasy® Blood and Tissue Kit (Qiagen, Venlo, Netherlands).

Tanks used for behavioral observations (36L x 33W x 24H cm) had lines drawn on the bottom to separate it into equal thirds (i.e., left, center, right). Each observation tank was lined with gravel, but the floor was cleared immediately around the lines so that they were still visible from a “top-down” view for the novel object test. A plastic plant was placed into the middle third of each tank to provide a refuge. Opaque dividers were inserted between the observation tanks during behavioral testing; otherwise individuals had visual access to fish in neighboring tanks.

***Measuring neophilia as response to a novel object.*** To set up the neophilia assay, the plastic plant was removed, plastic dividers were placed on all sides of the tank, and a mirror was positioned at a 45-degree angle above the tank to provide a top-down view. A camcorder (JVC Everio HD Hard Dish Camcorder Model No: GZ-HD40U) was used to record the trial via the mirror. A perforated tank divider was used to block the fish from accessing the back part of the tank that was out of the camera’s view. One of the outside thirds of the tank was randomly selected, and a circular blind was used to corral the individual into that area. The novel object was then placed in the opposite end. After five minutes, the blind was removed. The observer recorded how much time was spent orienting towards the novel object while within one body length of the object. Upon completion of the test, the home tank was reset to its pretrial state.

***Measuring flexibility with the barrier task.*** All barrier detour task-related trials occurred in a separate testing tank. This tank had the same dimensions as the individual’s home observation tank. No gravel was present to help increase the salience of the food reward. To begin the session, the individual was gently scooped with a white cup from their home tank and transferred to an opaque shelter that was then placed into the back-center of the testing tank and the fish was left undisturbed for three minutes. A 60W mm circle was initially drawn on the center of the floor of the tank (~7 cm from the entrance of the shelter), and a single blood worm was placed within that circle. Between trials fish were allowed to reacclimate to the shelter for two minutes. After the fourth trial of the day the fish were returned to their home tanks. If an individual did not reach criterion within four days, an additional individual from that family was sampled later in the experiment if possible, in order to maximize the number of individuals tested within each family. 15 individuals did not reach criterion. For the actual barrier trial fish were given 30 minutes to emerge and solve the task. Eleven fish did not solve the task and eat within 30 minutes. All fish approached and ate the food on the fourth trial that followed the barrier trial, suggesting that those that did not solve the task were still motivated by the food reward. Upon completion of the fourth trial, the individual was returned to its home tank.

***Compliance with ethical standards.*** Fish were caught in the field using baited minnow traps under the approval of the State of Alaska’s Department of Fish and Wildlife permit to M. Bensky (FRP P-17-019 and FTP 17A-0020). In the lab, the fish were housed in groups before and after the experiment. All experimental procedures were non-invasive. While the fish were undergoing training they were housed individually, but were given visual access to neighboring fish when they were not participating in active trials in order to enhance their welfare. Fish were transferred from their home tank to the training tanks by gently scooping them in a cup to minimize stress.

***Statistical analysis.*** When there was a significant effect of population, we first tested the hypothesis that marine and well-established freshwater populations differ by creating a contrast matrix that directly compared these two population types (Team 2013)(package = “multcomp”; function = “glht”). We then performed post-hoc tests to compare the newly established freshwater populations. Contrast matrices were constructed so that post-hoc p-values were adjusted for multiple comparisons. Post-hoc contrasts focused on the two newly derived populations (Cheney Lake and Loberg Lake) because visual inspection of the data suggested that they are different. The first post-hoc contrast was between Cheney Lake and Loberg Lake. Then Cheney Lake, the newest freshwater population (8 years from collection), was compared to the established freshwater populations, while Loberg Lake (28-34 years from collection) was compared to the marine fish, and Cheney Lake and Loberg Lake were directly compared.

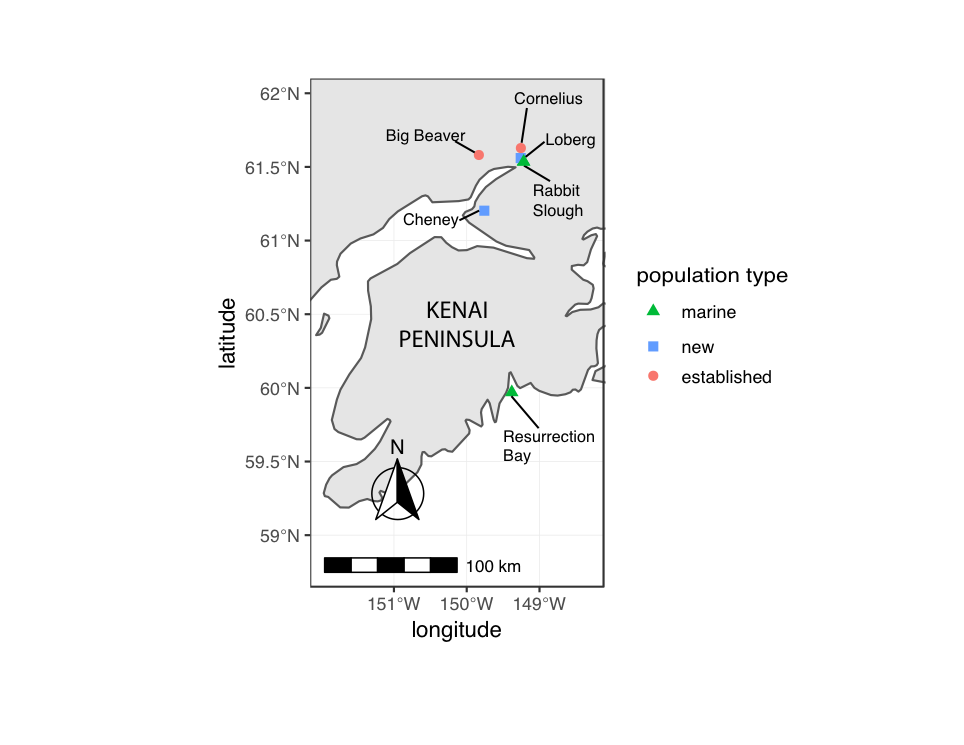

**Fig. S1.** Map of sampling sites.

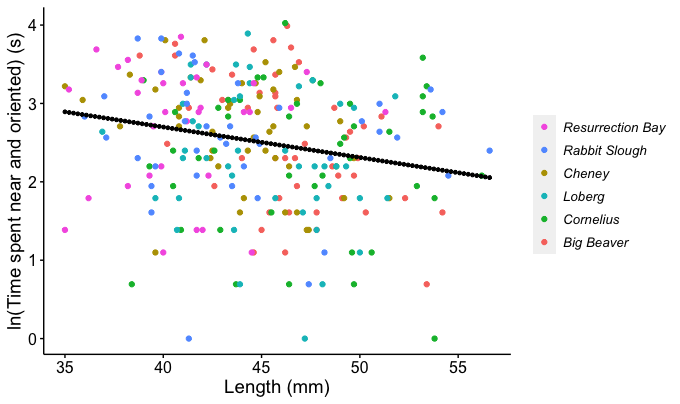

Fig. S2. Bigger fish were less neophilic (spent less time near and oriented to the novel object). Each data point represents a different individual, color coded by population.

**Table S1**. Linear mixed models testing the influence of population, length and sex on boldness (latency to emerge), flexibility (time at the apex) and neophilia (time near and oriented to the novel object. Significant terms are in bold.

| Behavior | Factor | Sum Sq | Mean Sum Sq | Num DF | Den DF | F | p-value |
| --- | --- | --- | --- | --- | --- | --- | --- |
| Boldness (latency to emerge) | | | | | | | |
|  | **Population** | **14.969** | **2.994** | **5** | **58.946** | **3.299** | **0.011** |
|  | Length | 0.037 | 0.037 | 1 | 253.774 | 0.041 | 0.840 |
|  | Sex | 2.610 | 2.609 | 1 | 248.498 | 2.875 | 0.091 |
| Flexibility (time at the apex) | | | | | | | |
|  | **Population** | **15.459** | **3.092** | **5** | **58.009** | **3.495** | **0.008** |
|  | Length | 0.0001 | 0.0004 | 1 | 237.932 | 0.001 | 0.982 |
|  | Sex | 1.008 | 1.008 | 1 | 235.742 | 1.140 | 0.287 |
| Neophilia (time near and oriented to the novel object) | | | | | | | |
|  | Population | 1.474 | 0.295 | 5 | 59.568 | 0.512 | 0.766 |
|  | **Length** | **3.986** | **3.986** | **1** | **247.793** | **6.923** | **0.009** |
|  | Sex | 0.407 | 0.407 | 1 | 253.628 | 0.707 | 0.401 |

**Table S2**. **Broad-sense heritabilities and genetic correlations within each population.** Heritabilities are on the diagonal, genetic correlations are on the off-diagonal, with 95% confidence intervals in parentheses. Bold indicates that the estimate is significantly different from zero. Greater boldness trait values correspond to lower boldness (i.e., slower to emerge), greater flexibility trait values correspond to less flexibility (i.e., more time persisting at the apex of the barrier)..

| **BIG BEAVER.** Sample size for correlations with flexibility is n=43, otherwise n=50 | | | |
| --- | --- | --- | --- |
|  | Boldness | Flexibility | Neophilia |
| Boldness | **0.754 (0.153, 1.000)** | -0.588 (-0.880, 0.295) | -0.219 (-0.747, 0.443) |
| Flexibility | -- | 0.225 (0.000, 0.974) | -0.327 (-0.782, 0.431) |
| Neophilia | -- | -- | 0.096 (0.000, 0.437) |
| **CHENEY LAKE.** Sample size for correlations with flexibility is n=43, otherwise n=46 | | | |
|  | Boldness | Flexibility | Neophilia |
| Boldness | 0.252 (0.000, 0.983) | -0.718 (-0.925, 0.134) | 0.171 (-0.584, 0.667) |
| Flexibility | -- | 0.325 (0.000, 0.994) | 0.0144 (-0.574, 0.647) |
| Neophilia | -- | -- | 0.151 (0.000, 0.684) |
| **CORNELIUS.** Sample size for correlations with flexibility is n=45, otherwise n=46 | | | |
|  | Boldness | Flexibility | Neophilia |
| Boldness | **0.737 (0.092, 1.000)** | **-0.777 (-0.931, -0.231)** | **-0.762 (-0.899, -0.206)** |
| Flexibility | -- | **0.864 (0.378, 1.000)** | **0.599 (0.053, 0.895)** |
| Neophilia | -- | -- | **0.876 (0.458, 1.000)** |
| **LOBERG LAKE**. Sample size for correlations with flexibility is n= 44, otherwise n= 45 | | | |
|  | Boldness | Flexibility | Neophilia |
| Boldness | 0.103 (0.000, 0.511) | -0.619 (-0.903, 0.282) | -0.586 (-0.862, 0.381) |
| Flexibility | -- | 0.692 (0.014, 1.000) | 0.585 (-0.258, 0.856) |
| Neophilia | -- | -- | 0.378 (0.000, 0.993) |
| **RABBIT SLOUGH.** Sample size for correlations with flexibility is n=40, otherwise n=42 | | | |
|  | Boldness | Flexibility | Neophilia |
| Boldness | 0.322 (0.000, 0.991) | -0.663 (-0.903, 0.298) | -0.206 (-0.691, 0.614) |
| Flexibility | -- | 0.329 (0.000, 0.996) | 0.094 (-0.571, 0.758) |
| Neophilia | -- | -- | 0.183 (0.000, 0.892) |
| **RESURRECTION BAY.** Sample size for correlations with flexibility is n=31, otherwise n=33 | | | |
|  | Boldness | Flexibility | Neophilia |
| Boldness | **0.700 (0.012, 1.000)** | -0.756 (-0.936, 0.259) | -0.237 (-0.746, 0.614) |
| Flexibility | -- | 0.279 (0.000, 0.995) | -0.011 (-0.700, 0.753) |
| Neophilia | -- | -- | 0.355 (0.000, 0.993) |

**Table S3**. **Phenotypic correlations (Pearson) within each population.** Bold indicates P<0.05.

| **ALL.** Sample size for correlations with flexibility is n=247, otherwise n=262 | | | |
| --- | --- | --- | --- |
|  | Boldness | Flexibility | Neophilia |
| Boldness | -- | **-0.522** | **-0.315** |
| Flexibility |  | -- | **0.229** |
| Neophilia |  |  | -- |
| **BIG BEAVER.** Sample size for correlations with flexibility is n=43, otherwise n=50 | | | |
|  | Boldness | Flexibility | Neophilia |
| Boldness | -- | **-0.297** | **-0.422** |
| Flexibility |  | -- | 0.038 |
| Neophilia |  |  | -- |
| **CHENEY LAKE.** Sample size for correlations with flexibility is n=43, otherwise n=46 | | | |
|  | Boldness | Flexibility | Neophilia |
| Boldness | -- | **-0.576** | -0.100 |
| Flexibility |  | -- | 0.156 |
| Neophilia |  |  | -- |
| **CORNELIUS.** Sample size for correlations with flexibility is n=45, otherwise n=46 | | | |
|  | Boldness | Flexibility | Neophilia |
| Boldness | -- | **-0.618** | **-0.554** |
| Flexibility |  | -- | **0.450** |
| Neophilia |  |  | -- |
| **LOBERG LAKE.** Sample size for correlations with flexibility is n=44, otherwise n=45 | | | |
|  | Boldness | Flexibility | Neophilia |
| Boldness | -- | **-0.402** | **-0.422** |
| Flexibility |  | -- | **0.293** |
| Neophilia |  |  | -- |
| **RABBIT SLOUGH.** Sample size for correlations with flexibility is n=40, otherwise n=42 | | | |
|  | Boldness | Flexibility | Neophilia |
| Boldness | -- | **-0.440** | -0.085 |
| Flexibility |  | -- | 0.228 |
| Neophilia |  |  | -- |
| **RESURRECTION BAY.** Sample size for correlations with flexibility is n=31, otherwise n=33 | | | |
|  | Boldness | Flexibility | Neophilia |
| Boldness | -- | **-0.478** | -0.127 |
| Flexibility |  | -- | 0.021 |
| Neophilia |  |  | -- |

**Table S4**. **Repeatability of latency to emerge across four trials.** Repeatability estimate was calculated using Bayesian statistics with Markov Chain Monte Carlo simulations using MCMCglmm package (Hadfield 2010) in R 3.5.3 (http://www.r-project.org/). A Poisson distribution was used because latency to emerge was positively skewed and used full integer time durations. We used non-informative proper priors with 1 000 000 iterations, thinning of 100 iterations, and burn-in of 100 000 iterations. Through these simulations, 95% confidence intervals were generated with significance of our estimate inferred according to whether the lower bound of the interval approached zero.

| **Population** | **n** | **Repeatability** | **Lower, Upper CI** |
| --- | --- | --- | --- |
| Big Beaver | 50 | 0.6985 | (0.5620,0.7866) |
| Cornelius | 46 | 0.6083 | (0.4782,0.7367) |
| Loberg Lake | 45 | 0.6565 | (0.4950,0.7553) |
| Cheney Lake | 46 | 0.6499 | (0.5385,0.7824) |
| Rabbit Slough | 42 | 0.5645 | (0.3961,0.7019) |
| Resurrection Bay | 33 | 0.7528 | (0.6218,0.8605) |

**Movie S1 (separate file):** Boldness assay (emergence from a refuge)

**Movie S2 (separate file)**: Flexibility assay (response to a barrier)
